# Rapid Decline Of Nesting Peregrine Falcons In The San Francisco Bay Region Of California Synchronous With An H5N1 Outbreak

**DOI:** 10.64898/2026.02.11.705416

**Authors:** Zeka E. Glucs, W. Grainger Hunt, A. Marm Kilpatrick, Jan Ambrosini, Daniel J. Armstrong, Douglas A. Bell, Jon Bianchi, Shirley Doell, Hannah Doniach, Benjamin M. Dudek, Nick Dunlop, Gavin Emmons, David Gregoire, Andre Gregoire, Alicia Lewis, John Lewis, Mary Malec, Stephanie Martin, Lori Michel, Denise F. Peck, Glenn R. Stewart, Tim Stroshane, Craig Ow, Bruce E. Lyon

## Abstract

After rebounding from near extirpation during the organochlorine era, breeding Peregrine Falcons (*Falco peregrinus*) in California are again facing adversity, this time consistent with an outbreak of a highly pathogenic avian influenza. Following the first detection of the H5N1 variant clade 2.3.4.4b virus in California wild birds in July 2022, we assembled data from long-term monitoring (2000–2025) of peregrine breeding territory occupancy in the broad vicinity of San Francisco Bay to examine possible impacts on falcon populations. Prior to the outbreak, 47 focal breeding territories had shown nearly complete occupancy by pairs (98.5% of 390 site-years), with very few vacancies, single birds in attendance, or subadult pair members. Within 8 mo of the outbreak, occupancy had dropped to 65.1%, and 2 yrs later (2025), only 36.2% of sites remained occupied. An uptick in site-occupancy by single birds also occurred after the outbreak, but it is unclear whether these were survivors or floaters attempting to fill vacant territories where both pair members had perished. The high vacancy rates signaled an impact upon floaters (nonbreeding adults) that normally buffer breeding site-occupancy in healthy peregrine populations. From October 2022 through November 2025, 17 peregrine fatalities were diagnosed with H5N1 within the counties comprising our study area. Evidence that H5N1 caused these territory vacancies includes, (1) the striking temporal coincidence of occupancy loss with the outbreak, and (2) the lethality of the virus to peregrines and its confirmed presence in peregrine prey in our study area. Our study reaffirms the value of long-term territory occupancy monitoring in this sentinel species.

## INTRODUCTION

The peregrine falcon (*Falco peregrinus*) is an iconic avian top-predator with a remarkable history during the latter half of the 20^th^ century. Bioaccumulation of DDE, a stable metabolite of the organochlorine insecticide DDT introduced to agriculture in 1947, caused peregrines to lay thin-shelled eggs that failed to hatch (Cade et al. 1997, White et al. 2024). By the 1960s, widespread reproductive failure had extirpated nesting peregrines across the Eastern US and caused major declines in the rest of North America (Hickey 1969). The number of pairs in California fell from well over 150 to an estimated maximum of five by 1970 (Bond 1946, Herman 1971). A breeding program, initiated by falconers and academics under the umbrella of The Peregrine Fund and the Predatory Bird Research Group, University of California, Santa Cruz, produced thousands of young peregrines for release to the wild (Cade and Burnham 2003). The program was so successful that peregrines were removed from the US Endangered Species list in 1999. Recovery continued apace after that point, with effective saturation of breeding territories over most of their former range in recent decades (White et al. 2024).

There is now increasing evidence that peregrines have suffered another demographic setback—this time from Highly Pathogenic Avian influenza (HPAI; Harvey et al. 2023, Badia-Boher et al. 2025, Lambertucci et al. 2025). Peregrines prey upon a wide variety of taxa known to be important HPAI reservoirs (White et al. 2024), including Charadriiformes (Fig. 1) and Anseriformes (Andreasen et al. 2025, Haman et al. 2024, Hill et al. 2022, Runstadler and Puryear 2024, Damodaran et al. 2025); peregrines are thus particularly likely to encounter this deadly virus.

**Figure 1.**
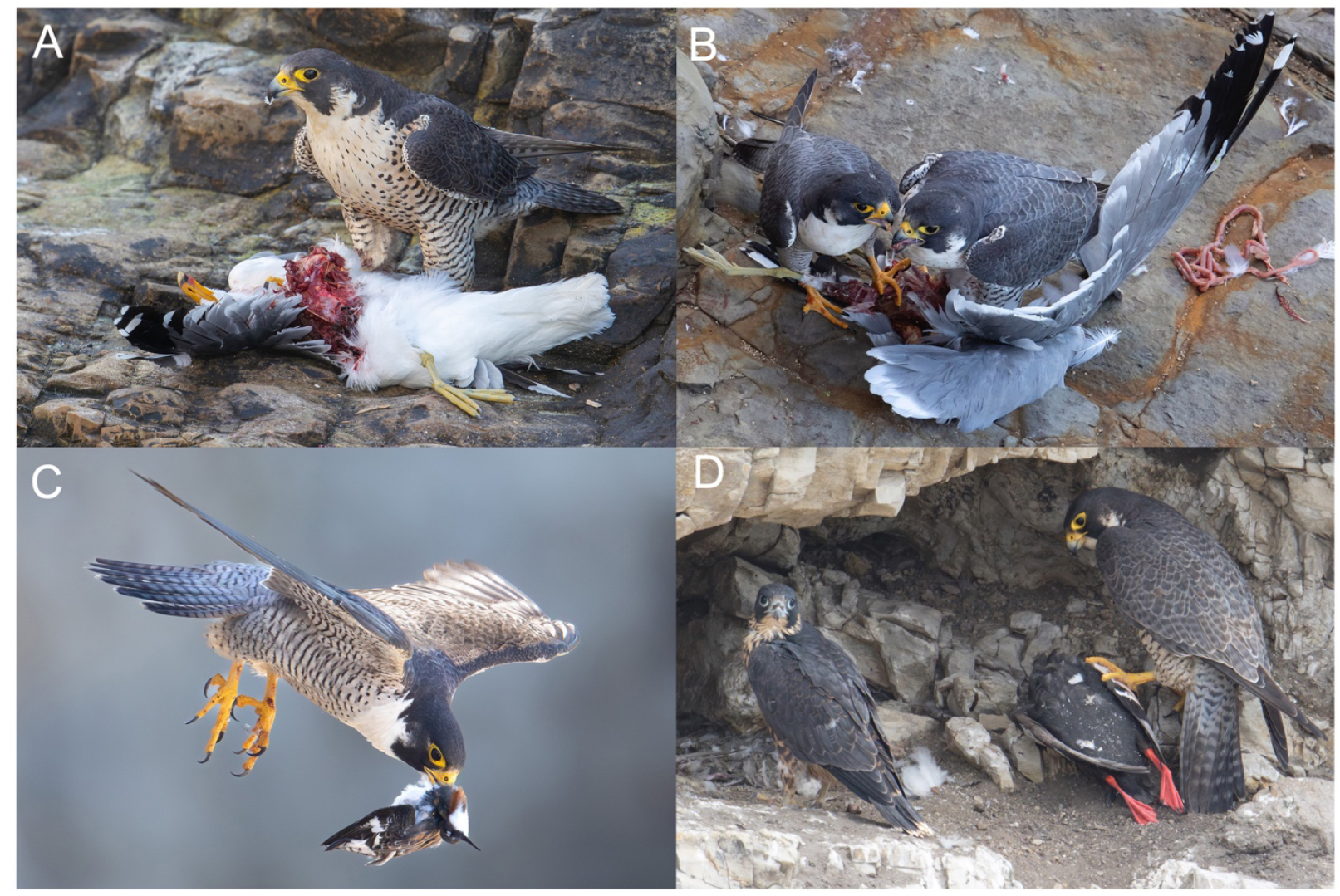
Nesting falcons from our study population consuming Charadriiform birds, important HPAI reservoir species: (A) California gull (*Larus californicus*), Laridae. (B) A breeding falcon pair sharing a California gull. (C) Red-necked phalarope (*Phalaropus lobatus*), Scolopacidae. (D) Pigeon guillemot (*Cepphus columba*), Alcidae. All photos Bruce Lyon.

Avian influenza is a widespread disease of wild birds, with a long evolutionary and ecological relationship. It has been of low virulence in chronically infected wild birds (Lycett et al. 2019), whereas lethal outbreaks are frequently documented in domesticated populations (Lycett et al. 2019, Lambertucci et al. 2025). In the last two decades, however, the transmission dynamics and virulence of the disease have changed dramatically (Atkinson and Baillie 2025, Caliendo et al. 2025). Spillover from domestic hosts back to wild birds is thought to be the driver of the change in dynamics (Ramey et al. 2022).

The HPAI variant, H5N1, was first detected in Guangdong, China, in 1996 but was restricted to domestic birds at the time (Klaassen and Willie 2023). An apparent key shift occurred in late 2002 to 2003, and significant numbers of fatalities were observed in several wild waterbird species in Hong Kong (Ramey et al. 2018). Since then, the H5N1 variant has spread to most continents, affecting numerous species, and the temporal and spatial patterns of prevalence and severity of outbreaks have changed (Caliendo et al. 2022). Due to this dramatic increase in geographic and host range, H5N1 was considered an ‘emerging disease threat’ to wild bird populations (Ramey et al. 2022).

Two significant HPAI outbreaks have occurred in North American wild birds. Several H5 strains were detected in late 2014 (Ip et al. 2015), lingering until 2016 when they abruptly disappeared (Krause et al. 2016). Then, in 2020, a massive global outbreak of H5N1 variant clade 2.3.4.4b (hereafter H5N1) occurred and caused what has been claimed as the most severe panzootic ever recorded (Lambertucci et al. 2025). Mass mortality events have been reported for both birds and mammals around the globe, with infections documented in over 400 bird species and 50 mammal species (Lambertucci et al. 2025). Evidence from several review studies suggest that some raptor species may be particularly vulnerable, especially scavengers and bird-eaters (Nemeth et al. 2023, Hall et al. 2024, Ringenberg et al. 2024). Given the low population densities of raptors and the consequent low probability of encountering their fatalities, demographic studies rather than simple counts of infected or dead birds are needed to more fully determine population impacts of this virus.

We here report a sudden reduction in breeding territory occupancy within a long-studied peregrine population in the region surrounding San Francisco Bay; the decline occurred during the H5N1 outbreak first detected in California in mid-2022 (California Department of Fish and Wildlife 2022, US Department of Agriculture 2025). Guiding our assessment are features of peregrine life history and population dynamics that underlie the known stability of healthy populations (Ratcliffe 1993). Pairs of this highly territorial species are widely spaced, requiring cliffs (or artificial structures) with specific characteristics appropriate to survival and nest success (White et al. 2024). Healthy populations tend toward territory saturation after which substantial numbers of floaters (adults without territories) accumulate and then stabilize (Hunt 1998). Floaters fill breeding vacancies as they occur, thus maintaining adult occupancy of virtually all territories. With moderate decreases in survival or reproduction, the floating population may dwindle to the point at which yearling pair members (subadults) are able to fill territory vacancies (see Tordoff and Redig 1997, Hunt and Law 2023). With more extreme impacts upon vital rates, territories may contain only single birds or become vacant. Two recent studies of peregrines in Europe illustrate these demographic effects in the presence of H5N1. Smith et al. (2025) observed an increase of breeding falcons with immature plumage in Scotland in 2023 but did not find a significant reduction in territory occupancy. In contrast, Badia-Boher et al. (2025) documented a sudden 25% reduction in occupancy in a population in the Netherlands over a two-year period.

Our study addresses several questions relating to the most recent H5N1 outbreak. Broadly, did it have detectable effects on our study population? Was the outbreak followed by a decrease in breeding territory occupancy, as occurred in the Netherlands (Badia-Boher et al. 2025)? Or were the effects more subtle, with only an increase in subadult pair members as observed in Scotland by Smith et al. (2025)? To explore whether something else could explain changes in occupancy, we assessed reproduction prior to the outbreak. We also compared these data to reproductive success post-outbreak to determine whether this rate changed. We examined the patterns specific to urban-versus rural-nesting birds, the two subpopulations having been differentiated in a previous demographic analysis (Kauffman et al. 2003) and an analysis of nesting habitat (Venu 2018). Finally, we contrasted changes in occupancy at inland sites versus coastal and bay sites, the latter having more immediate access to waterbirds known to be reservoirs of H5N1.

## METHODS

### Study Area and Data Collection

Since the Peregrine Falcon’s delisting in 1999, the Predatory Bird Research Group has monitored a resident coastal nesting population from the Monterey Bay region north to Point Reyes and the associated coastal and mid-coastal mountain ranges. The latter consist of rolling hills supporting redwood forests, and with oak woodland further east. The Sacramento River estuary at the heart of the region contains tidal marshes and a diversity of waterbirds. San Francisco Bay itself is rimmed by cities and urban sprawl; certain buildings, bridges, and other artifacts offer nesting places for peregrine pairs, and scattered cliffs in the rural environment provide many additional sites. The climate is Mediterranean, with dry summers and wet winters. Our study area covers about 8,000 km^2^ of the roughly 11000-km^2^ area collectively known as the Greater San Francisco Bay Area.

In 2008, the Predatory Bird Research Group began a formal banding program in the study area aimed at detecting fatalities, natal dispersal, and breeder turnover, with an additional goal of encouraging community members and university students to observe falcon nesting territories and search for new ones. Observations took place annually during January–July and were initially biased towards nests where property owners or managers granted access for banding young. In 2018, our monitoring goals shifted towards surveying all known nesting territories in the study area (50–60 sites annually).

Observers visited each territory at least twice per breeding season between March and June; each was recorded as occupied by a pair or a single individual, or judged vacant (Fig. 2). Making that determination sometimes required three or more hours of observation or repeated visits, depending on the stage of the breeding cycle and other factors. We were also alert to the presence of subadult (yearling) pair members, based on plumage (White et al. 2024). Once a site was deemed occupied, we returned periodically to determine reproductive outcome. We quantified reproduction as the average number of large young observed (≥ 34 days post-hatch) per pair and fledging success as the proportion of nests containing large young in a given year. Nest cameras at five nest sites functioned as gateways to public appreciation and concern for the species and produced valuable details relating to the current study.

**Figure 2.**
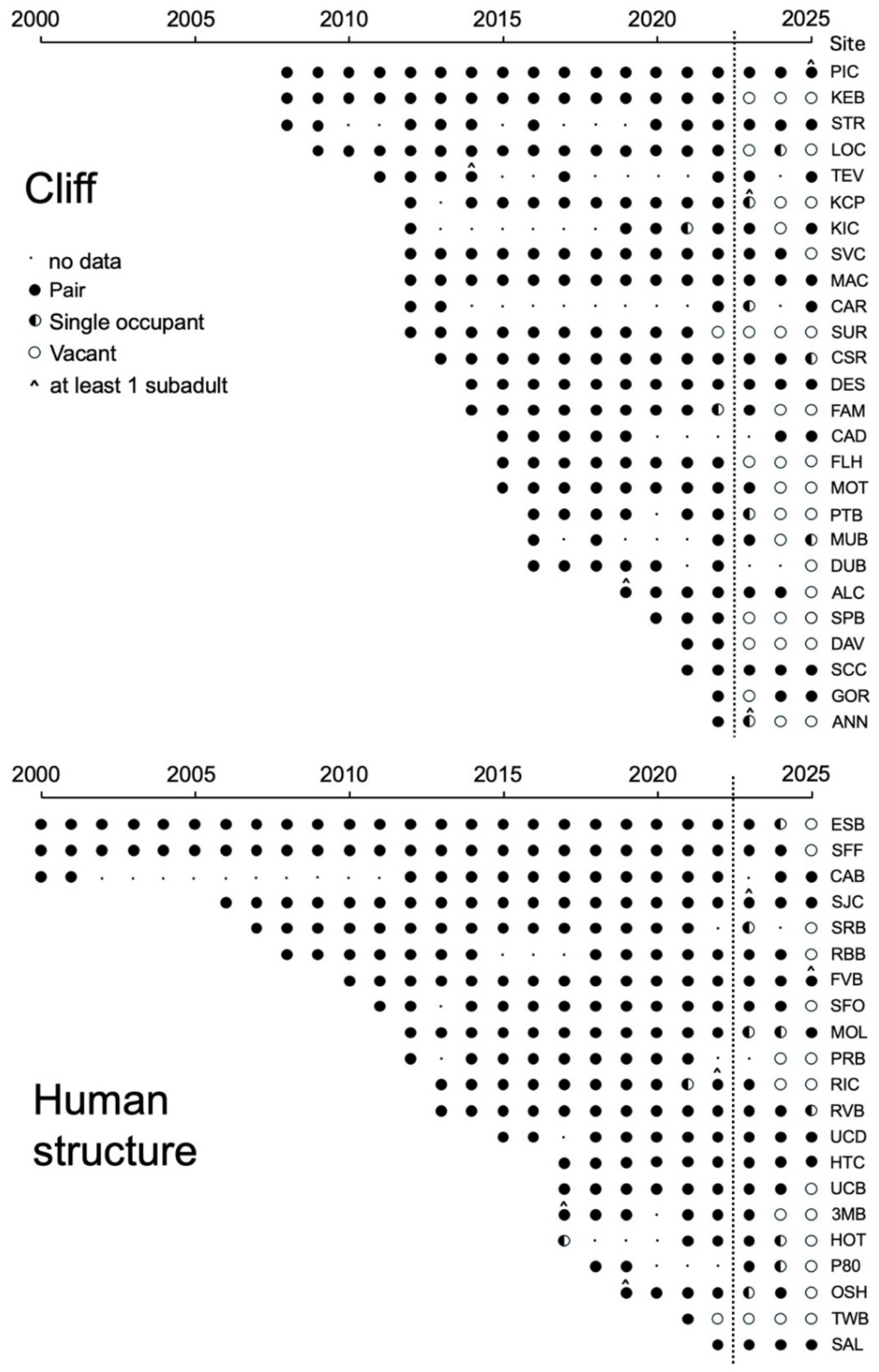
Timelines of occupancy patterns of individual sites before and after the outbreak of H5N1 in California in 2022 (dashed vertical line). Previous work suggested distinct demography for urban versus rural sites, so patterns are shown separately. The increasing number of sites over time reflects both increased sampling effort and population growth.

### Criteria for Site Inclusion in this Study

H5N1 variant clade 2.3.4.4b was first found in North America in late 2021, but it was not detected in California wild birds until July 2022 (California Department of Fish and Wildlife 2022, US Department of Agriculture 2025); we used the latter date for separating our observations into pre- and post-outbreak, with spring 2023 being the first post-outbreak breeding season for peregrines in our study area. To examine changes in peregrine demography before and after the H5N1 arrival, we chose breeding sites that (1) had been censused both before and after the outbreak, (2) had produced at least one clutch of eggs since territory establishment, and (3) were known to have been occupied in 2021 or 2022, or, in the case of sites not surveyed in either of those years, we included those occupied in the previous two years the site was surveyed. Forty-seven sites met these criteria and are included in our analyses. Thirteen were excluded, three of which did not meet criterion 1; seven failed criterion 2, and three failed criterion 3. Not all sites were checked in all years (2000– 2025), and only in 2025 did we check every site in this dataset.

### Statistical Approach

We used General Linear Mixed Models (GLMM) with a binomial distribution (and logit link) to compare pair occupancy, the proportion of territories with single birds, and the proportion of territories with subadult birds before and after the outbreak using the *lme4* package in R (version 4.3.3). Two time-periods (before, after the outbreak) were included as a fixed effect, and site was included as a random effect. We conducted two additional analyses with GLMM to determine if pair occupancy differed across time periods for (1) birds nesting on human structures (a proxy for urban nesting) and those using cliffs and (2) sites on the coast or San Franscisco Bay compared to inland sites (the former having access to waterbirds known to be H5N1 reservoirs).

## RESULTS

### Site Occupancy Changes after Outbreak

Prior to the H5N1outbreak in California in mid-2022, the 47 Peregrine Falcon breeding sites we monitored were almost aways occupied by pairs (Fig. 2, 3). Of 390 site-years monitored between 2000 and 2022, 384 (98.5%, 95% CI: 96.7% – 99.4%) contained a pair, 4 had a single bird (1.0%, 95% CI: 0.28% – 2.6%), and 2 (0.5%, 95% CI: 0.062% – 1.8%) were unoccupied (Figs. 2, 3). In 2023, the year after H5N1 was first detected in California, only 28 of 43 territories (65.1%, 95% CI: 49.1% – 79.0%) were occupied by pairs (Figs. 2, 3A). The next year, 2024, pair occupancy declined to 51% (22 of 43, 95% CI: 35.5% – 66.7%), and by 2025, it dropped to 36.2% (17 of 47, 95% CI: 22.7% – 51.5%) (Figs. 2, 3A). Overall, the proportion of territories occupied by pairs was significantly lower after the 2022 outbreak than before (GLMM with a binomial distribution and territory ID as a random effect: post-2022 coeff.: -4.87 ± SE 0.57, Z = 8.5, P < 2×10^-16^). The rates of decline (slopes in Fig. 3) were similar across the 3 years following the outbreak: a 30% reduction in sites occupied by pairs from 2022 to 2023, then a 21% reduction in 2024, and a 23% reduction in 2025. Interestingly, we recorded no fully unoccupied sites until spring 2022 when two vacancies were documented (Fig. 2); these observations occurred in the months before the outbreak was detected (July 2002), suggesting the possibility of an earlier onset of the epidemic.

**Figure 3.**
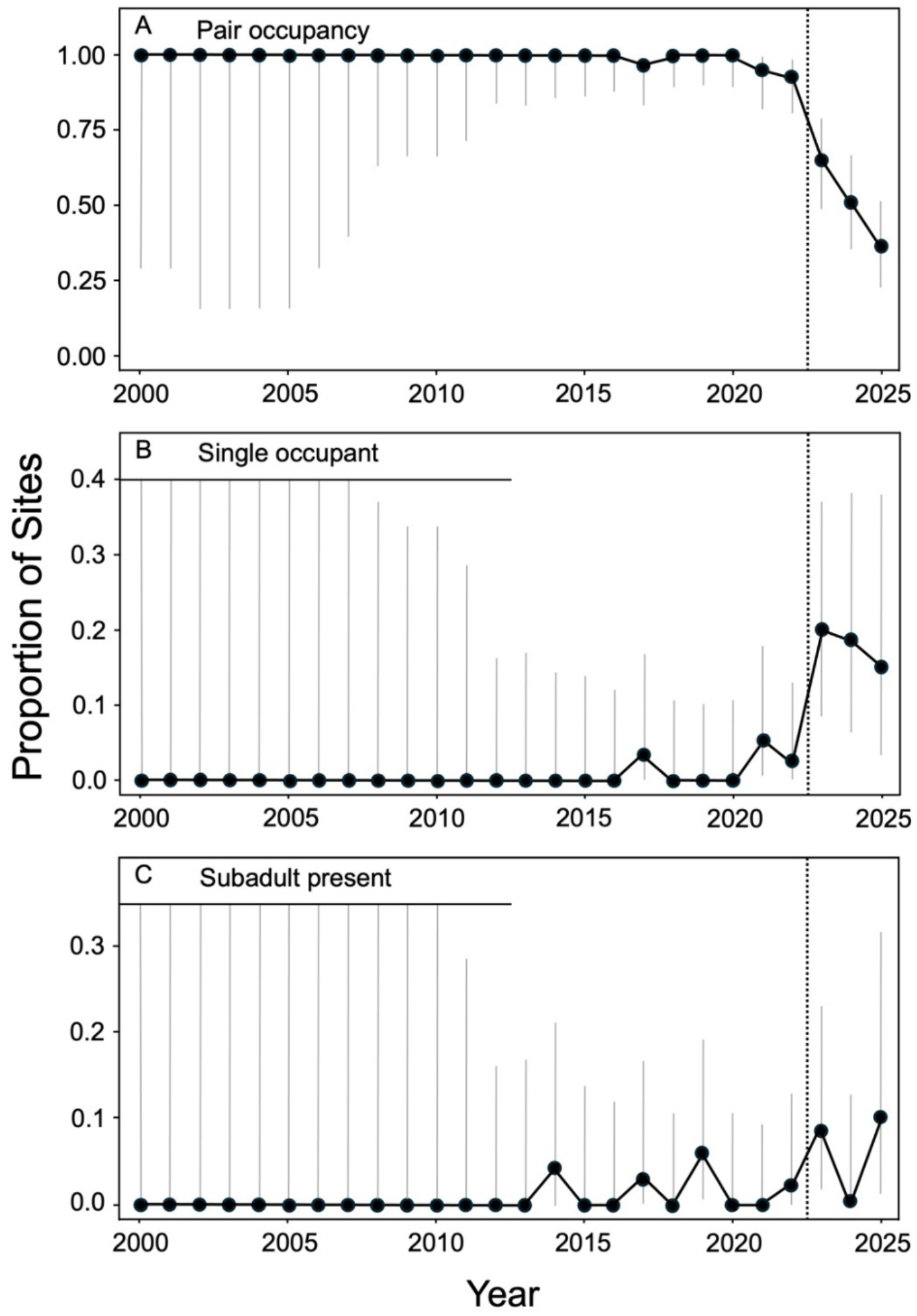
Population summary of the occupancy patterns shown in Figure 2. Fig. 3A shows occupancy by pairs among all territories surveyed in each specific year; Figs. 3B and 3C mark the proportion of single occupants and subadults, respectively, among tenanted sites. Vertical dashed line marks the first detection of H5N1 in California wild birds (July 2022); error bars show 95% confidence intervals; upper limits are truncated in early years at thin horizonal line in panels B and C.

Sites with single birds present or subadult pair members can indicate mortality and possible deficiency in the floater buffer (see Hunt and Law 2003), so we examined whether their frequencies changed after the outbreak. For this analysis, we excluded vacant sites and focused on sites with at least one bird. There was a sharp increase in the percentage of sites occupied by a single bird immediately following the outbreak (Figs. 2, 3B; GLMM with a binomial distribution and territory ID as a random effect: post-2022 coeff.: 3.10 ± SE 0.60, Z = 5.16, P = 2.44 × 10^-7^). Prior to the outbreak, single birds were detected in only four (1.0%, 95% CI: 0.28% – 2.6%) of 388 site-years with birds present; in the three years following the outbreak (2023–2025), single attendants occupied 20.0% (7/28), 18.5% (5/27), and 15.0% (3/20) of the non-vacant sites, respectively (Figs. 2, 3B). The frequency of subadult birds attending territories also increased after the outbreak (Figs. 2, 3C; GLMM with a binomial distribution and territory ID as a random effect: post-2022 coeff.: 1.60 ± SE 0.64, Z = 2.49, P = 0.013). Five of the 388 (1.3%, 95% CI: 0.42% –3.0%) site-years with at least one bird prior to the outbreak had a subadult bird present whereas following the outbreak, a subadult was present in 5 of 82 (6.1%; 2.0% – 13.7%) site-years where at least one bird was present.

Visually comparing the locations of sites that became vacant after the outbreak with those that remained occupied did not reveal clear spatial patterns (Fig. 4). We found no difference in pair occupancy decline between territories with cliff nests and those with nests on human structures (GLMM with a binomial distribution, with pre-post-2022 interacting with territory type as fixed effects and territory ID as a random effect, post-2022 coeff.: -5.27± SE 0.80, Z = 6.61, P = 4.0 × 10^-11^; human structure coeff.: 0.60± SE 0.62, Z = 0.97, P = 0.33; interaction term (human-post) coeff.: 0.84 ± SE 1.01, Z = -0.82, P = 0.41). For example, in 2025, 38% (10 of 26) cliff sites were occupied compared to 33% (7 of 21) nests associated with human structures (Fig. 2). Finally, we compared changes in pair occupancy of territories in coastal/bay versus inland areas and found no difference (coastal/bay pair occupancy 2023– 2025: 50/95, 52.6% (95% CI: 42.1% – 63.0%); inland: 17/38, 44.7% (95% CI: 28.6% - 61.7%); GLMM with a binomial distribution, with pre-post-2022 interacting with region as fixed effects and territory ID as a random effect, post-2022 coeff.: -4.55± SE 0.87, Z = 5.23, P = 1.67 × 10^-7^; coastal coeff.: 0.45 ± SE 0.67, Z = 0.67, P = 0.50; interaction term (post-2022 – coastal): coeff.: - 0.49 ± SE 1.04, Z = 0.47, P = 0.64).

**Figure 4.**
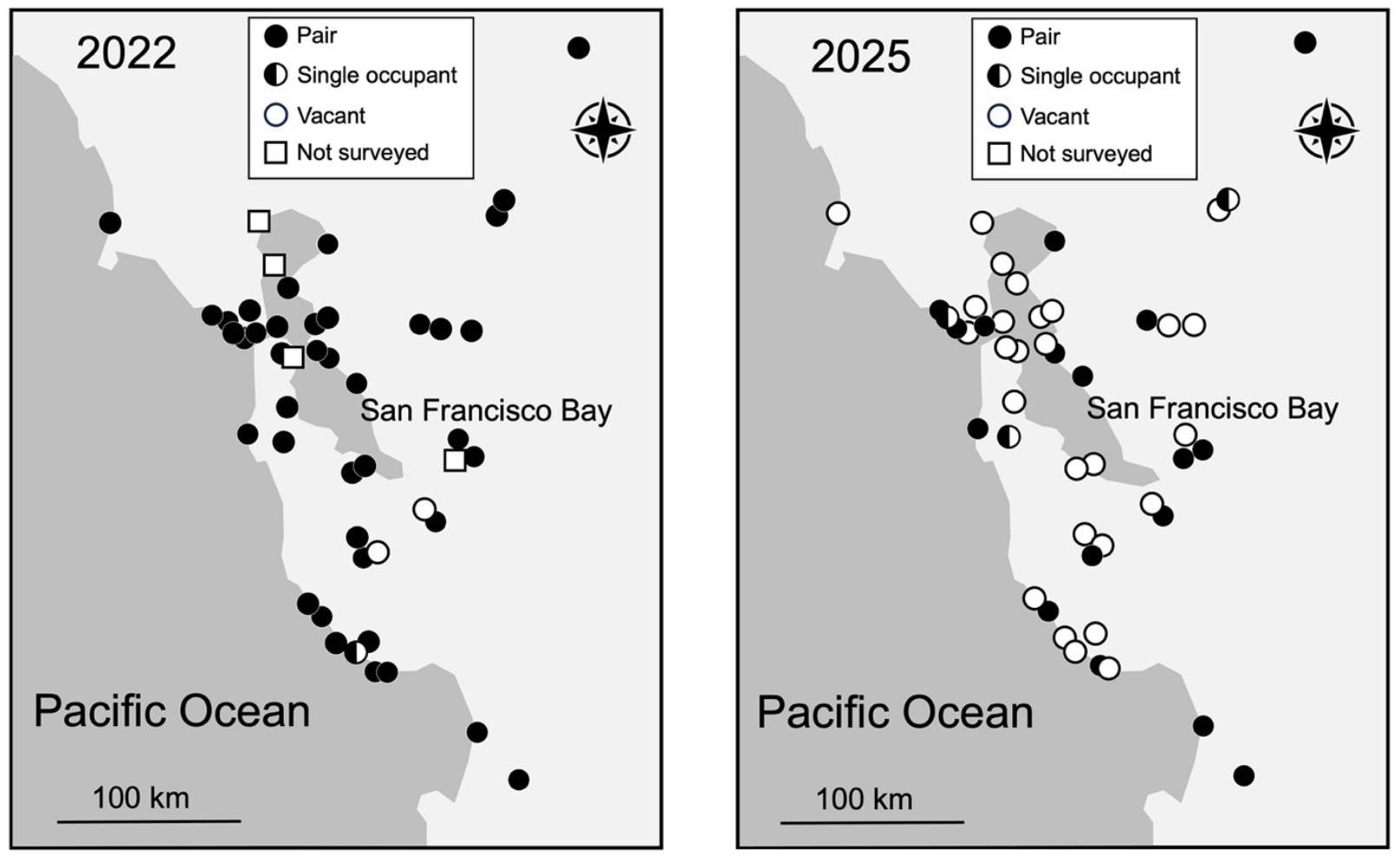
Contrasting spatial patterns of site occupancy for two years: 2022, the breeding season immediately before the outbreak was detected in California, and 2025, the most recent territory census.

**Figure 5.**
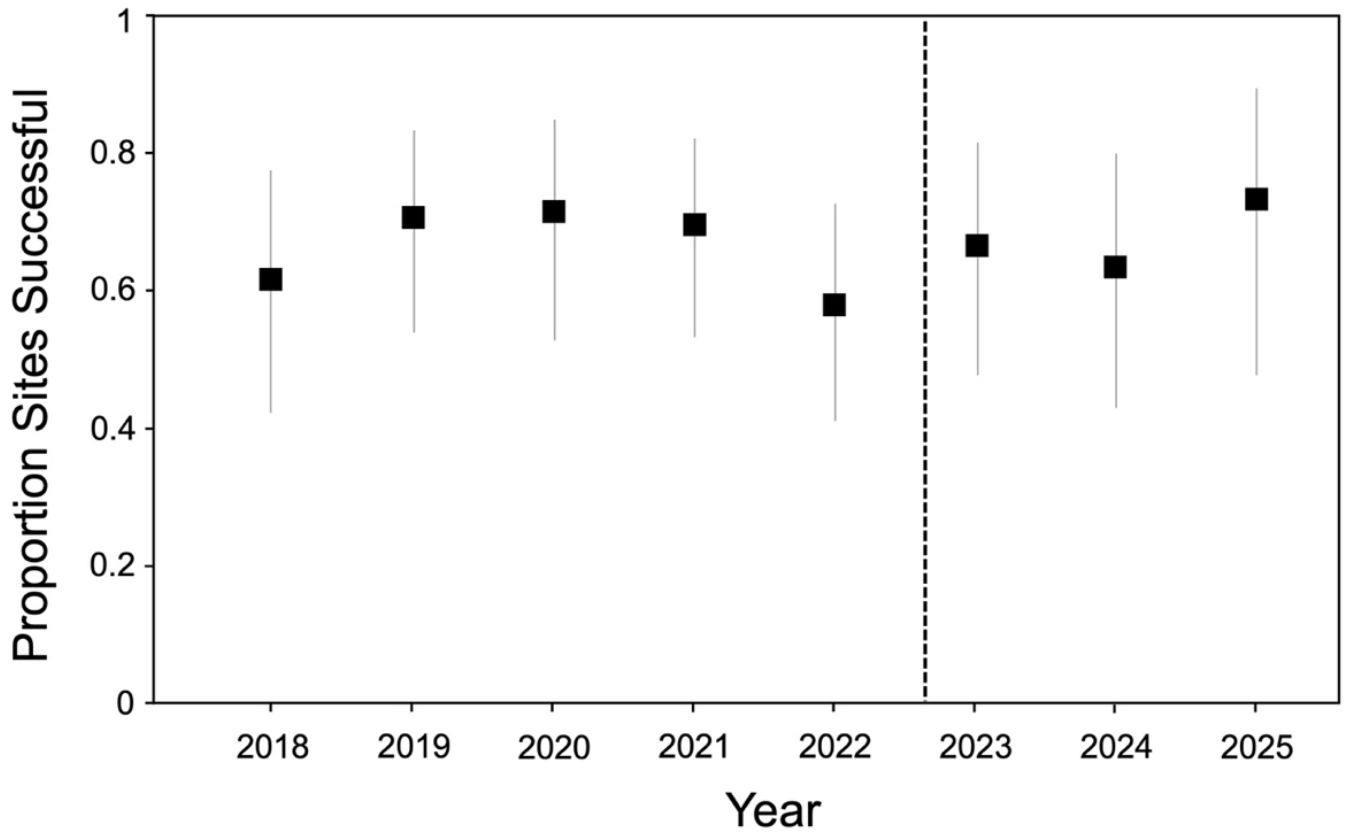
Fledging success (fledged young vs. unsuccessful) of breeding territories from 2018 to 2025. Dashed vertical line separates the periods before and after the outbreak. Symbols show proportion of sites that fledged at least one young; vertical lines are 95% confidence intervals.

### Fledging Success

To determine whether nest failure was associated with the outbreak, we compared the proportions of site-years that fledged young. Fledging success was virtually identical before (2018–2022) and after (2023–2025) the outbreak: 66.5% (105 of 158 site-years; 95% (CI: 58.5% – 73.8%) produced fledglings before the outbreak compared to 67.2% (43 of 64 site-years after the outbreak; 95% CI: 54.3% – 78.4%) (GLMM with a binomial distribution, with territory ID as a random effect, post-2022 coeff.: - 0.28± SE 0.38, Z = 0.72, P = 0.47). We also assessed whether there were indications of reproductive problems prior to the outbreak on the sites that would later become vacant. Limiting the period to before the outbreak, the proportion of site-years that each site produced young did not differ between sites that were not occupied by a pair in 2025 (68 of 108; 63.0%, 95% CI: 53.1% – 72.1%) and those that were occupied by a pair (37 of 50; 74%, 95% CI: 59.7% – 85.4%; GLMM with a binomial distribution, with territory ID as a random effect, 2025 pair occupancy coeff.: 0.84 ± SE 0.67, Z = 1.26, P = 0.21).

## DISCUSSION

Prior to the H5N1 clade 2.3.4.4b outbreak, our sample population of 47 peregrine territories appeared robust. Nesting territories were almost invariably occupied by pairs (98.5% of site-years), exhibited a normal reproductive rate (see below), and only rarely contained a single occupant or subadult pair member (Figs. 2, 3). By all expectations, breeding habitat was at or near saturation, and floaters were quickly replacing missing occupants and contesting territory ownership. Then in spring 2023, 8 mos after the first detection of H5N1 in wild birds in California (July 2022), our annual survey of nesting peregrines in the study area revealed a sudden substantial reduction in territory occupancy. By 2025, just 2 yrs later, only 36% of the original 47 territories still contained pairs (Figs. 2, 3); the frequency of single occupants had also increased, as did that of subadult pair members.

Whereas the dramatic increase in breeding territory vacancies we document is clearly dire, it does not account for an additional impact upon the floater population. We used simple equations from Hunt (1998) and basic survival and reproductive data for our population to crudely estimate this component of Moffat’s equilibrium. Previous work in our study area produced estimates of annual survival at 0.86 for adults and 0.29 for juveniles (Kauffman et al. 2003). With these rates and an average of 1.7 fledglings per occupied site (ZEG, unpubl. data), some 38% of the adult population would have been floaters capable of replacing dead breeders. Floater presence almost certainly underpinned the consistency we observed in site-occupancy prior to the outbreak and may explain the resilience of a Scottish population that suffered mortality from H5N1 (Smith et al. 2025). A presumed contraction of the floater population in our study area may at least partly account for the absence of occupancy renewals apparent in Fig. 2.

Our data may also underestimate mortality among territory-holders in that some sites that remained occupied after 2022 may have contained replacement pair members. Notably, the banded breeding female of a closely monitored pair in the city of San Jose was observed on camera showing clear neurological symptoms on 26 February 2023 just prior to the breeding season. She ultimately fell from her perch, was recovered the next morning, then died in captivity the evening of 27 February. Her mate, also banded, was last seen on 28 February. A new, unbanded adult female appeared in the nest box on 3 March, and an unbanded subadult male arrived shortly thereafter. The two have since held the territory and consistently reproduced, even in the year they arrived.

Evidence that H5N1 caused the breeding vacancies we report is mostly circumstantial, but corroborating details relate to its presence in the ecosystem. The exact timing of the disappearances with respect to the onset of the outbreak is strongly inferential, together with known lethality to peregrines of those strains of H5N1 then prevalent in California wild birds. The US Department of Agriculture (USDA) reported 26 H5N1 infections in wild peregrine fatalities collected in California during October 2022–November 2025 (Appendix 1). Likewise, 72 dead or dying gulls of five species (mostly *Larus occidentalis* and *L. californicus*) tested positive for H5N1 in California during that same period (US Department of Agriculture 2025). The nine counties containing our study area accounted for 17 (65%) of the 26 H5N1 peregrine fatalities reported in California, and 39% of gulls (an indicator of H5N1 prevalence). In all, the US Department of Agriculture reported over 1,400 H5N1-diagnosed fatalities among wild birds in California through November 2025. Surveys elsewhere have found very few peregrines with seroprevalence of antibodies indicating they contracted and survived the disease; only 2% of 205 falcons sampled in migration had antibodies for influenza A viruses, which included H5N1 (Earthspan Foundation Org 2025).

Gulls, shorebirds, alcids, and waterfowl are frequent prey of peregrines in our study area, and all are known reservoirs of H5N1 (Hill et al. 2022, Haman et al. 2024, Runstadler and Puryear 2024, Andreasen et al. 2025, Damodaran et al. 2025). Peregrines, with their extraordinarily diverse diet, are particularly susceptible to viral contact with these and other species, numbering in hundreds of individuals per year per individual falcon. Given the neurological effects of H5N1, birds infected with the disease would be more quickly overtaken by their pursuers, peregrines being known to exploit disadvantaged prey and even carrion, factors compounding their susceptibility to infection (White et al. 2024, Caliendo et al. 2025). In a wintering population along coastal Washington state, peregrines scavenged 30% of their food items, and waterbirds were most likely to be scavenged (Varland et al. 2018).

The preponderance of abrupt territory vacancies rather than a more gradual stepwise attrition of individual territory-holders appears novel for this species. The explanation might be that a pair member contracting H5N1 prior to or during the breeding season may infect its mate either directly or by sharing prey (Fig. 1B). Both members may thus contract and succumb to the virus, and the territory may lie vacant until a prospecting floater settles there.

While our evidence that H5N1 caused excessive mortality among territory-holders and floaters within our study area is substantial, reproduction was apparently uncompromised at sites that remained occupied by pairs post-outbreak. Our finding of no change in reproductive rate during the H5N1 outbreak is similar to that observed by Badia-Boher et al. 2025 in the Netherlands; interestingly, both areas contain extensive tidal wetlands and wintering migratory waterbirds. The lack of impact on reproduction might be explained by an apparent reduction in viral presence in spring; 14 (87%) of the 17 peregrine fatalities with H5N1 diagnoses (2022-2024) reported by the US Department of Agriculture in the Greater San Francisco Bay counties took place during October–February (Supplemental Table S1); the remaining three were collected in March during which most pairs in the region begin incubating. Winter-diagnosed H5N1 gull fatalities, as indications of viral presence, did not extend past 6 March.

A question remains as to the geographic scale and patchiness of the decline we have documented. Public reports from Yosemite National Park in the Sierra Nevada Mountains have indicated no apparent loss in occupancy or reproduction among its 16 monitored pairs (National Park Service 2025). Perhaps peregrines nesting elsewhere in the mountainous interior have been similarly less affected. If true, some degree of floater buffering and repatriation from such environments might be expected from migration into our study area, although this will likely be ineffective until the disease wanes.

Peregrine Falcons are again showing themselves as indicators of environmental perturbation. As before, when knowledge of the norms of territory occupancy and diet led to an understanding of biomagnification by a harmful contaminant (DDE), the species now marks the severity of a virulent pathogen within its food web. Our study thus reaffirms the value of long-term occupancy monitoring and collaboration among those who routinely monitor their local territories. We urge wider such cooperation across differing ecological settings, with focus on known, reliably censused territories where pre-outbreak data exist. Such information may be valuable in ascertaining the prospects of peregrines in the larger landscape during this period of uncertainty.

## SUPPLEMENTAL MATERIAL

(available online). Table S1. List of H5N1-diagnosed Peregrine Falcon fatalities in California through November 2025, as reported by the US Department of Agriculture.

## ACKNOWLEDGEMENTS

We acknowledge the recovery work of the Peregrine Fund and the Santa Cruz Predatory Bird Research Group for ensuring there are nesting peregrine falcons to observe in the wild. Countless volunteers and students have contributed hours of observational data to make this study possible. Collection of nest data in the mosaic of land use within the San Francisco and Monterey Bay areas has required the cooperation of many landowners and agencies, including: National Park Service, California State Parks, CalTrans, East Bay Regional Parks, Santa Clara County Parks, Midpeninsula Regional Open Space District, San Francisco Public Utilities Commission, San Jose City Hall, University of California, Bay Area Raptor Rescue, and several private companies. This manuscript was improved by comments from Bob Montgomerie and editing by Terry Hunt. We thank the Ahmanson Foundation and the Helen and Will Webster Foundation for funding.

This research was completed under the following permits and approvals: CAFWS Scientific Collection Permit SC-010757 and associated Peregrine Falcon MOU, USGS Bird Banding Permits 22383 and 24347, and University of California, Santa Cruz IACUC protocols Glucz2403dn and Glucsz2404dn.

---

**Supplemental Table S1.** List of H5N1-diagnosed Peregrine Falcon fatalities in California through November 2025, as reported by the US Department of Agriculture. Data downloaded from https://www.aphis.usda.gov/livestock-poultry-disease/avian/avian-influenza/hpai-detections/wild-birds

| State | County | In study area | Collection Date | Date Detected | HPAI Strain | Bird Species | WOAH Classification | Sampling Method | Submitting Agency |
| --- | --- | --- | --- | --- | --- | --- | --- | --- | --- |
| California | Alameda | Y | 2/14/24 | 3/4/24 | EA/AM H5N1 | Peregrine falcon | Wild bird | Morbidity/Mortality | CA DFW/CAHFS |
| California | Contra Costa | Y | 12/5/23 | 2/2/24 | EA H5N1 | Peregrine falcon | Wild bird | Morbidity/Mortality | CA DFW/CAHFS |
| California | Contra Costa | Y | 12/11/24 | 6/27/25 | EA/AM H5N1 | Peregrine falcon | Wild bird | Morbidity/Mortality | CA DFW/CAHFS |
| California | Del Norte | N | 12/7/22 | 12/21/22 | EA/AM H5N1 | Peregrine falcon | Wild bird | Morbidity/Mortality | CA DFW/CAHFS |
| California | Inyo | N | 11/29/23 | 12/13/23 | EA/AM H5N1 | Peregrine falcon | Wild bird | Morbidity/Mortality | Private (non-government) submission |
| California | Marin | Y | 12/1/23 | 2/2/24 | EA H5N1 | Peregrine falcon | Wild bird | Morbidity/Mortality | CA DFW/CAHFS |
| California | Merced | N | 11/13/23 | 9/19/24 | EA/AM H5N1 | Peregrine falcon | Wild bird | Morbidity/Mortality | CA DFW/CAHFS |
| California | Monterey | Y | 11/16/22 | 12/13/22 | EA/AM H5N1 | Peregrine falcon | Wild bird | Morbidity/Mortality | CA DFW/CAHFS |
| California | Napa | N | 10/3/22 | 10/19/22 | EA/AM H5N1 | Peregrine falcon | Wild bird | Morbidity/Mortality | CA DFW/CAHFS |
| California | Orange | N | 10/18/22 | 10/31/22 | EA/AM H5N1 | Peregrine falcon | Wild bird | Morbidity/Mortality | CA DFW/CAHFS |
| California | Sacramento | Y | 10/30/23 | 8/1/24 | EA/AM H5N1 | Peregrine falcon | Wild bird | Morbidity/Mortality | CA DFW/CAHFS |
| California | Sacramento | Y | 11/5/25 | 11/20/25 | EA/AM H5N1 | Peregrine falcon | Wild bird | Morbidity/Mortality | CA DFW/CAHFS |
| California | Sacramento | Y | 11/20/25 | 12/1/25 | EA H5 | Peregrine falcon | Wild bird | Morbidity/Mortality | CA DFW/CAHFS |
| California | San Diego | N | 11/22/22 | 12/13/22 | EA/AM H5N1 | Peregrine falcon | Wild bird | Morbidity/Mortality | CA DFW/CAHFS |
| California | San Diego | N | 11/1/23 | 11/14/23 | EA/AM H5N1 | Peregrine falcon | Wild bird | Morbidity/Mortality | CA DFW/CAHFS |
| California | San Luis Obispo | N | 11/16/22 | 12/13/22 | EA/AM H5N1 | Peregrine falcon | Wild bird | Morbidity/Mortality | CA DFW/CAHFS |
| California | San Luis Obispo | N | 1/24/23 | 2/10/23 | EA H5N1 | Peregrine falcon | Wild bird | Morbidity/Mortality | CA DFW/CAHFS |
| California | San Mateo | Y | 1/10/23 | 1/26/23 | EA/AM H5N1 | Peregrine falcon | Wild bird | Morbidity/Mortality | CA DFW/CAHFS |
| California | San Mateo | Y | 3/30/23 | 4/13/23 | EA/AM H5N1 | Peregrine falcon | Wild bird | Morbidity/Mortality | CA DFW/CAHFS |
| California | San Mateo | Y | 1/11/24 | 1/30/24 | EA/AM H5N1 | Peregrine falcon | Wild bird | Morbidity/Mortality | CA DFW/CAHFS |
| California | San Mateo | Y | 1/11/24 | 1/30/24 | EA/AM H5N1 | Peregrine falcon | Wild bird | Morbidity/Mortality | CA DFW/CAHFS |
| California | Santa Clara | Y | 3/16/23 | 4/12/23 | EA/AM H5N1 | Peregrine falcon | Wild bird | Morbidity/Mortality | CA DFW/CAHFS |
| California | Santa Clara | Y | 12/12/23 | 2/2/24 | EA H5N1 | Peregrine falcon | Wild bird | Morbidity/Mortality | CA DFW/CAHFS |
| California | Santa Clara | Y | 12/19/23 | 9/19/24 | EA/AM H5N1 | Peregrine falcon | Wild bird | Morbidity/Mortality | CA DFW/CAHFS |
| California | Santa Clara | Y | 3/26/25 | 5/28/25 | EA/AM H5N1 | Peregrine falcon | Wild bird | Morbidity/Mortality | CA DFW/CAHFS |
| California | Solano | Y | 2/27/25 | 6/17/25 | EA/AM H5N1 | Peregrine falcon | Wild bird | Morbidity/Mortality | CA DFW/CAHFS |

